# Dynamic variations of resting-state BOLD signal spectra in white matter

**DOI:** 10.1101/2021.07.15.452567

**Authors:** Muwei Li, Yurui Gao, Adam W. Anderson, Zhaohua Ding, John C. Gore

## Abstract

Recent studies have demonstrated that the mathematical model used for analyzing and interpreting fMRI data in gray matter (GM) is inappropriate for detecting or describing blood-oxygenation-level-dependent (BOLD) signals in white matter (WM). In particular the hemodynamic response function (HRF) which serves as the regressor in general linear models is different in WM compared to GM. We recently reported measurements of the frequency contents of resting-state time courses in WM that showed distinct power spectra which depended on local structural-vascular-functional associations. In addition, multiple studies of GM have revealed how functional connectivity between regions, as measured by the correlation between BOLD time series, varies dynamically over time. We therefore investigated whether and how BOLD signals from WM in a resting state varied over time. We measured voxel-wise spectrograms, which reflect the time-varying spectral patterns of WM time courses. The results suggest that the spectral patterns are non-stationary but could be categorized into five modes that recurred over time. These modes showed distinct spatial distributions of their occurrences and durations, and the distributions were highly consistent across individuals. In addition, one of the modes exhibited a strong coupling of its occurrence between GM and WM across individuals, and two communities of WM voxels were identified according to the hierarchical structures of transitions among modes. Moreover, the total number of transitions in each community predicts specific human behaviors. Last, these modes are coupled to the shape of instantaneous HRFs. Our findings extend previous studies and reveal the non-stationary nature of spectral patterns of BOLD signals over time, providing a spatial-temporal-frequency characterization of resting-state signals in WM.

## Introduction

Functional magnetic resonance imaging (fMRI) has been widely exploited for mapping brain activities by measuring changes in blood-oxygenation-level-dependent (BOLD) signals that are coupled with energy consumption by neurons^1^. Mathematical models have been developed to identify image voxels that exhibit increased signal amplitudes triggered by a task performed by a participant during a scan, thereby mapping the brain activation associated with the task^2^. In addition, BOLD signal fluctuations measured during a resting state reflect spontaneous variations in neural activities and reveal intrinsic functional networks characterized according to the degree of temporal synchronization among brain regions^3^. While reports of BOLD signals in gray matter (GM) have dominated the fMRI literature, there have been far fewer descriptions of BOLD effects in white matter (WM), partly because they are weaker (due to the smaller blood flow/volume compared to GM^4^), but also because of the use of inappropriate hemodynamic response functions which have been used as regressors to detect activations. However, a growing body of evidence suggests that BOLD signals can be reliably detected in WM and reflect neural activities^5,6,15,16,7–14^, and such signals are altered significantly in patients with neurological or psychiatric disorders^17–20^. Recent studies have demonstrated how the temporal profiles of WM BOLD responses to stimuli are different from GM, with reduced magnitudes and delayed peaks, reflecting differences in hemodynamic conditions between GM and WM^21–24^. These studies highlight the importance of understanding the characteristics of WM timecourses so that they can be incorporated appropriately into detection models.

We recently showed that the power spectra of WM time courses differ in shape from those in GM in the low-frequency (0.01-0.08 Hz) band, and they vary as a function of location and depend on the local neurovascular and anatomical configurations of WM^25^. Moreover, the spectral patterns at different locations predict their engagements in functional integration as well as specific human behaviors. These findings suggest the frequency contents of WM signals are heterogeneous spatially. However, potential temporal variations in BOLD signal characteristics in WM have not been investigated, reflecting an assumption that BOLD signals are statistically stationary. Here we examine this assumption and demonstrate that BOLD power spectra vary during a scan and reflect an average of multiple sub-patterns that recur over time.

A short-time Fourier transform (STFT) method was used to estimate the time-windowed frequency content of BOLD time courses from each WM voxel. A power spectrum was then calculated window-by-window, revealing the variations in amplitudes of frequencies with time. Our data suggest that the time-averaged spectrum (as reported in our previous study) is, in fact, the combination of five sub-patterns, which we call spectral modes (or frequency modes^26^), that recur over time. They include four unimodal modes (each of which exhibits single peaks at different frequencies, modes 1-4) and one uniform mode (with power distributed uniformly across the frequency band of interest at a relatively low level, mode 5). These modes show distinct spatial distributions of their occurrences and durations across WM voxels, and these distributions were highly consistent across individuals. One of the modes (mode 1) exhibited a strong coupling of its occurrence between GM and WM across individuals. More interestingly, two communities of WM voxels were identified according to the hierarchical structures of transitions among modes. Specifically, in community 1, the probability of transitions among modes 1, 2, and 5 are significantly higher than in community 2. By contrast, in community 2, the probability of transitions among modes 3, 4, and 5 are significantly higher than in community 1. Moreover, the total number of transitions in each community can predict specific human behaviors. Last, the instantaneous spectral patterns are associated with distinct HRFs that also vary with time. Our findings extend our previous study by looking at dynamic changes in BOLD spectra and reveal the non-stationary nature of spectral patterns as well as their physiological coupling over time, thereby providing a spatial-temporal-frequency characterization of resting-state BOLD signals in WM.

## Results

### Spectrogram of WM time courses

The spectrogram regarding each WM voxel was calculated using STFT methods. Briefly, each time course was split into 266 partially overlapped windows. Each window is 100.8 s in length, with a 97.92 s section overlapped with the previous window. In other words, the spectrogram provides a temporal resolution of around 3 s (2.88 s) and a frequency resolution down to around 0.01Hz. Figure 1 (middle column) shows examples of calculated spectrograms in which the selected WM voxels exhibit a single peak or dual peaks in their power spectra, similar to what we observed in our previous study. There is also another group of voxels that exhibit a flat, low-amplitude spectrum across frequencies. We also observed that the peaks of spectral patterns (Figure 1 middle column) are primarily determined by widely separated bursts of higher powers, as shown in Figure 1 left column.

**Figure 1.**
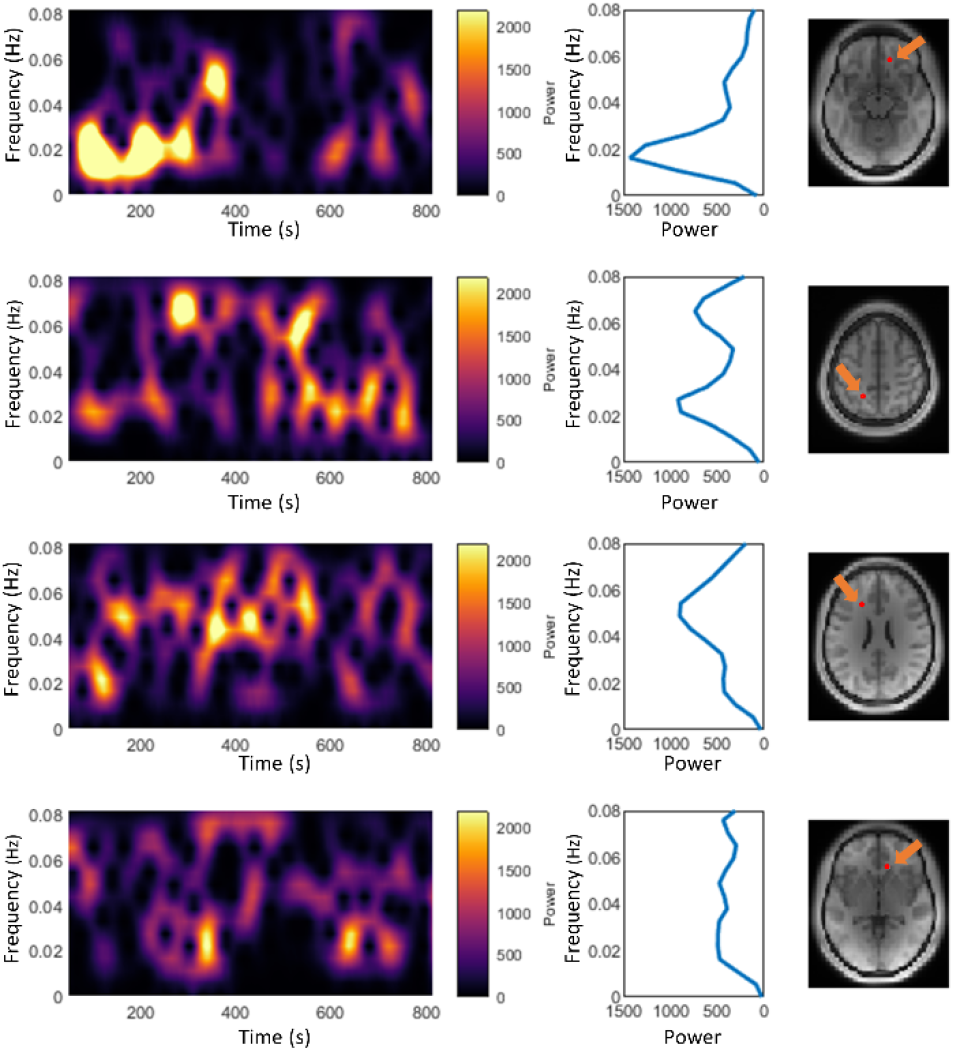
Examples of spectrograms of selected WM voxels in a single subject. Each row corresponds to one voxel whose spectrogram is shown on the left. The middle figure shows the mean power across all time windows, approximating the stationary power spectra of the signal. The red dot on the right figure indicates the location of the voxel.

### The spectral modes

As illustrated in Figure 2, five spectral modes were derived using k-means clustering of windowed spectral patterns (observations) across time windows, WM voxels, and subjects. Modes 1-4, which we called unimodal modes, exhibit single sharp peaks at different frequencies, while mode 5, is a uniform mode, with spectral power relatively constant and low across the frequency band of interest. As shown in Figure S1, the same workflow yielded two modes in GM, including one unimodal mode whose pattern is similar to mode 1 in WM and one uniform mode, which approximates to mode 5 in WM.

**Figure 2.**
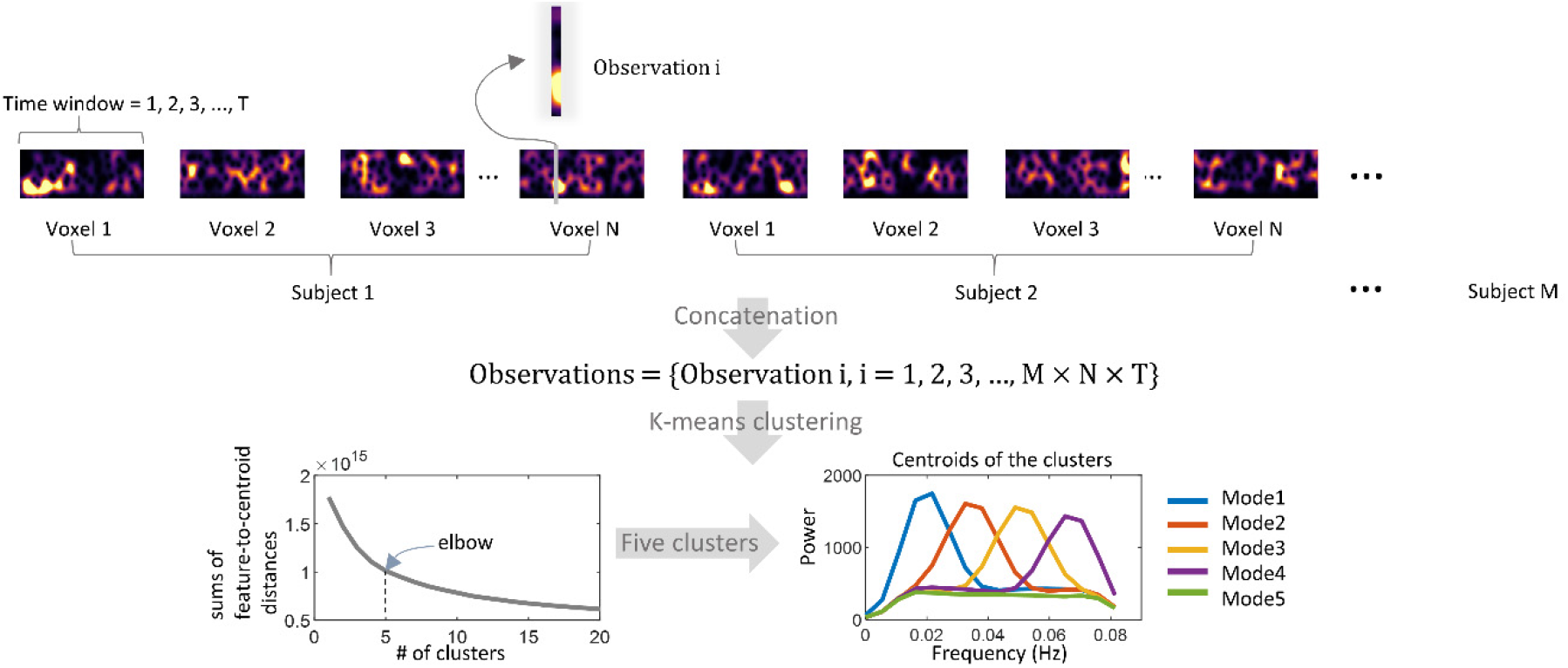
The workflow of calculating the spectral modes. A set of observations are generated by concatenating the spectrogram over voxels and subjects. Each observation is a vector, recording the spectra pattern of the signal corresponding to a specific time window. Observations are grouped into five clusters using the K-means method. The elbow criterium is used to determine the optimal value of the # of clusters (5 in the current study).

### Occurrence and duration of the modes

An occurrence map represents how often a mode occurs in every voxel during the entire scanning time. The distribution of voxels that exhibit significantly high occurrence of each mode, as visualized in 3D in Figure 3, was obtained by applying a one-sample t-test (p<0.05, FWE corrected) to the occurrence maps of all subjects. Figure S2 shows details of such distributions in axial slices, where the inferior frontal WM voxels exhibit significantly higher occurrences of modes 1 and 5 than other voxels. Meanwhile, the paraventricular and temporal voxels exhibit significantly higher occurrences of modes 3 and 4 than other voxels. However, the occurrences of mode 2 show little group effects, and so are not shown in Figure 3. In each panel of Figure 3, the boxplots represent the group-level distributions of the number of occurrences and the durations of a mode before a transition to another, measured from the area visualized in the brain maps shown to their left. We observed that mode 5 occurs most often and persists longest across all areas shown in Figure 3. In addition, in general, modes 1 and 5 occur more often and persist longer than modes 3 and 5 in the inferior frontal area, whereas they arise less often and last for shorter times in paraventricular and temporal areas, where modes 3 and 4 occur more often. Overall, the number of occurrences and durations is highly consistent across subjects, reflected by the tight interquartile range (distance between the upper bound and lower bound) of the boxes.

**Figure 3.**
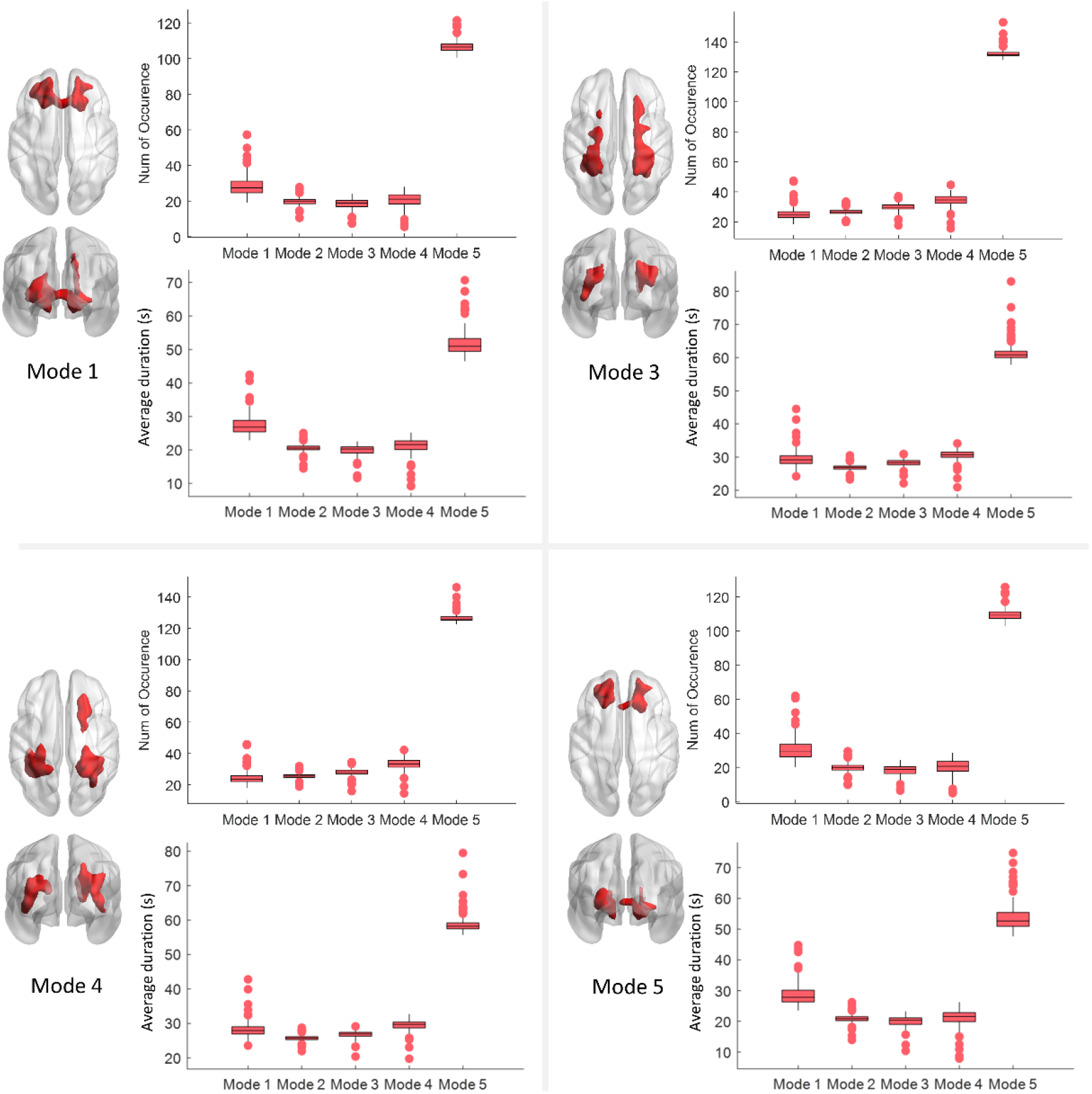
Voxels that exhibit significantly high occurrence of each mode across all subjects. In each of the four panels, the figure on the left visualizes the voxels that exhibit significantly higher occurrence of a certain mode than the rest of the WM voxels across all subjects (one-sample t-test, p<0.05 FWE corrected). The figure on the upper right of each panel shows the number of occurrences of the five modes within the area shown on the left figure. The figure on the lower right of each panel shows the average duration of the five modes within the area shown on the left figure. Each box visualizes the distribution of the measurements from all subjects. Note that the data for mode 2 is not shown because only a few voxels were found showing the significantly higher occurrence of mode 2 comparing to the rest of the WM voxels across subjects.

The distributions of voxels that exhibit significantly higher occurrences of the two modes in GM are shown in Figure S3. Cortical areas that are closer to the surface of the brain exhibited significantly higher occurrences of mode 1 than the rest of GM. Meanwhile, the voxels at insular, temporal areas and cingulate gyrus exhibit significantly higher occurrences of mode 2 than other GM voxels.

### Coupling of occurrence of modes between GM and WM

Figure 4 maps the coupling of the patterns of occurrences of specific modes that are prevalent in specific GM and WM voxels. The intensity of a WM voxel in the left figure indicates the Pearson’s correlation coefficient (r) between the occurrence of mode 1 at the voxel and the average occurrence of mode 1 in the GM region shown in Figure S3 (left) across subjects. It can be observed that a high occurrence of mode 1 in selected surface cortical areas reliably predicts high occurrences of mode 1 in multiple WM voxels in the same subject, and vice versa. The corresponding data that exceed a significant level (p<0.05, FWE corrected) are shown in Figure S4. For comparison, the coupling of occurrences of modes 2,3,4,5 in WM voxels to GM (mode 1) across subjects is shown in Figure S5.

**Figure 4.**
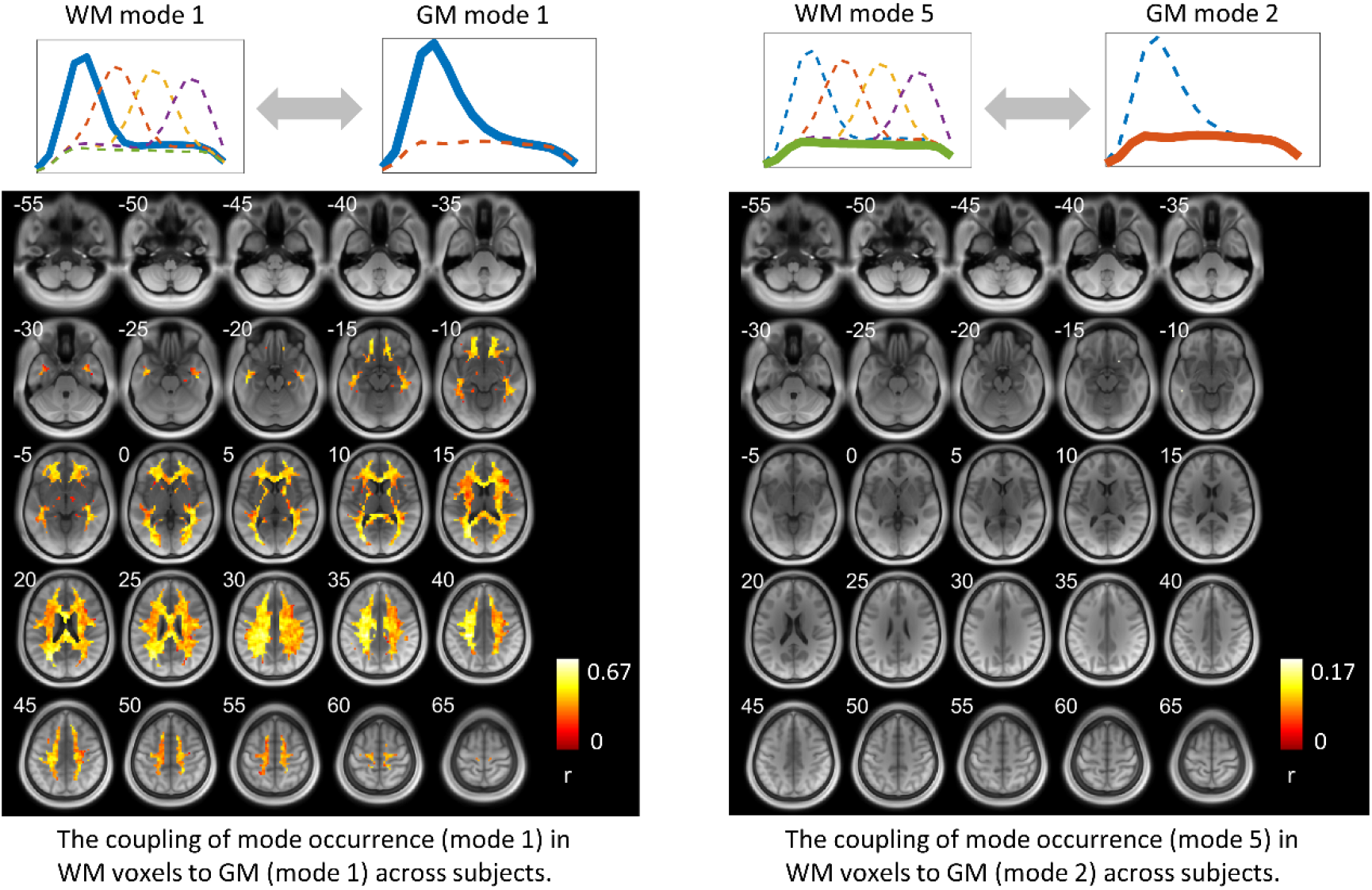
The coupling of mode occurrence in WM voxels to GM across subjects. The intensity of a voxel in the left figure indicates the Pearson’s correlation coefficient (r) between the occurrence of mode 1 at this WM voxel and the occurrence of mode 1 in the GM area shown in Figure S3 (left) across subjects. The intensity of a voxel in the right figure indicates the Pearson’s correlation coefficient (r) between the occurrence of mode 5 at this WM voxel and the occurrence of mode 1 in the GM area shown in Figure S3 (right) across subjects. Note that only significant (p<0.05) correlations are shown here.

The intensity of a voxel in the right figure of Figure 4 indicates the Pearson’s correlation coefficient (r) between the occurrence of mode 5 at the voxel and the average occurrence of mode 2 in the GM region shown in Figure S3 (right) across subjects. Only a few WM voxels show significant couplings of the occurrence of mode 5 to that of mode 2 in the GM region.

### Transitions among the modes

Figure 5 maps the number of transitions (shown as T values) among the five modes for every WM voxel. T values that are beyond a significant level (p<0.05, FWE corrected) are visualized in Figure S6. It can be observed that the inferior frontal area exhibits a significantly low number of transitions compared to the rest of WM.

**Figure 5.**
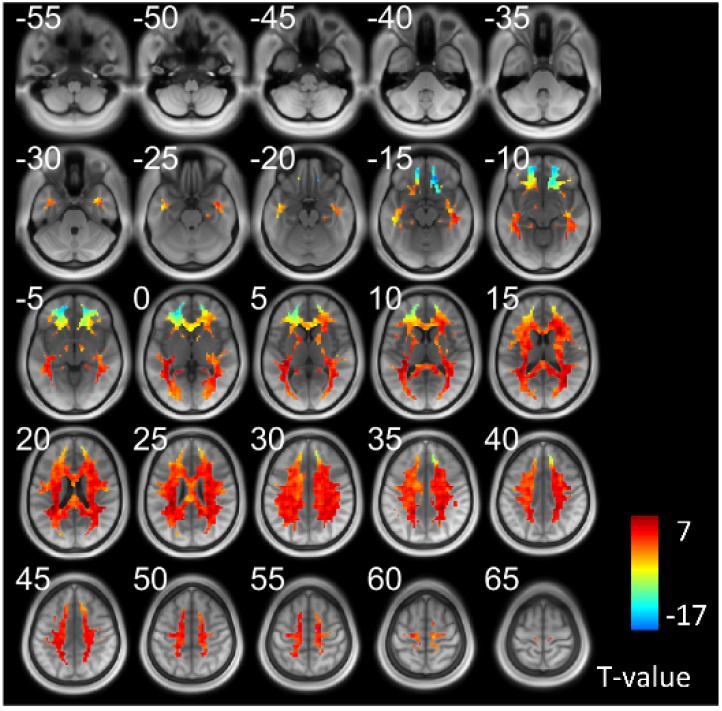
The number of transitions at each WM voxel across all subjects. T value was generated by a one-sample t-test across all subjects. Voxels that exabit significantly high/low values can be found in supplementary file.

Figure 6 shows the voxels that exhibit a significantly higher transition from mode i to mode j than the rest of WM voxels across subjects (one-sample t-test, p<0.05, FWE corrected). Two communities of voxels could be clearly identified and are coded by different colors. The lower left panel shows the voxels that exhibit significantly higher transitions among modes 1, 2, and 5 (community 1), along with its transition probability among the five modes shown on the right. The lower right panel shows the voxels that exhibit significantly higher transitions among modes 3, 4, and 5 (community 2), along with its transition probability among the five modes shown on the right. Note that the map with respect to each community was produced by a one-sample t-test (p<0.05, FWE corrected) on the transition maps among modes 1, 2, 5 or among modes 3, 4, 5 across all subjects. We observed that the patterns of transition probabilities are, in general, consistent between the two communities. However, a careful inspection reveals that community 1 exhibits a higher transition probability among modes 1, 2, 5, whereas a lower transition probability among modes 3, 4, 5 than community 2. Note that we have hidden the self transitions, which occupy more than 90% of the entire transitions.

**Figure 6.**
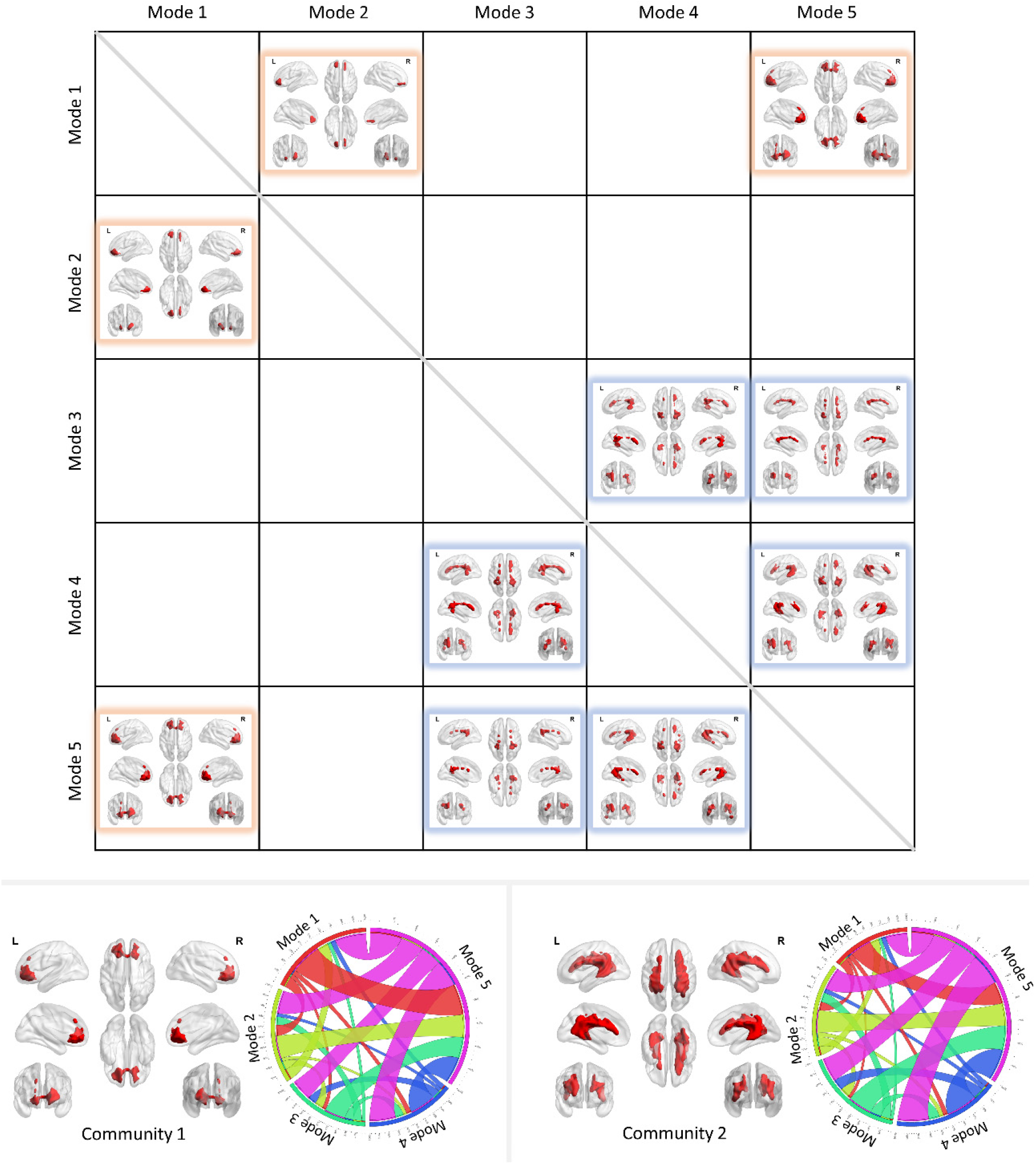
The transitions among modes. The upper figure visualizes the voxels that exhibit significantly higher transitions from mode i to mode j than the rest of WM voxels across subjects (one-sample t-test, p<0.05, FWE corrected). Two communities were identified and coded by different colors. The lower left panel shows the voxels that exhibit significantly higher transitions among modes 1, 2 and 5 (community 1), along with its transition probability among five modes shown on the right. The lower right panel shows the voxels that exhibit significantly higher transitions among modes 3, 4 and 5 (community 2), along with its transition probability among the five modes shown on the right.

### The correlation between modes and behavioral scores

Figure 7 shows the significant correlations between the number of transitions in each of the two communities and behavioral scores. We observed that more than half of the scores that are significantly predicted are associated with working memory functions. The number of transitions in community 2 shows more negative correlations than community 1, mostly associated with response times (RT) in working memory tasks. The significant correlations between the occurrence of the modes in each of the two communities and behavioral scores are shown in Figure S7.

**Figure 7.**
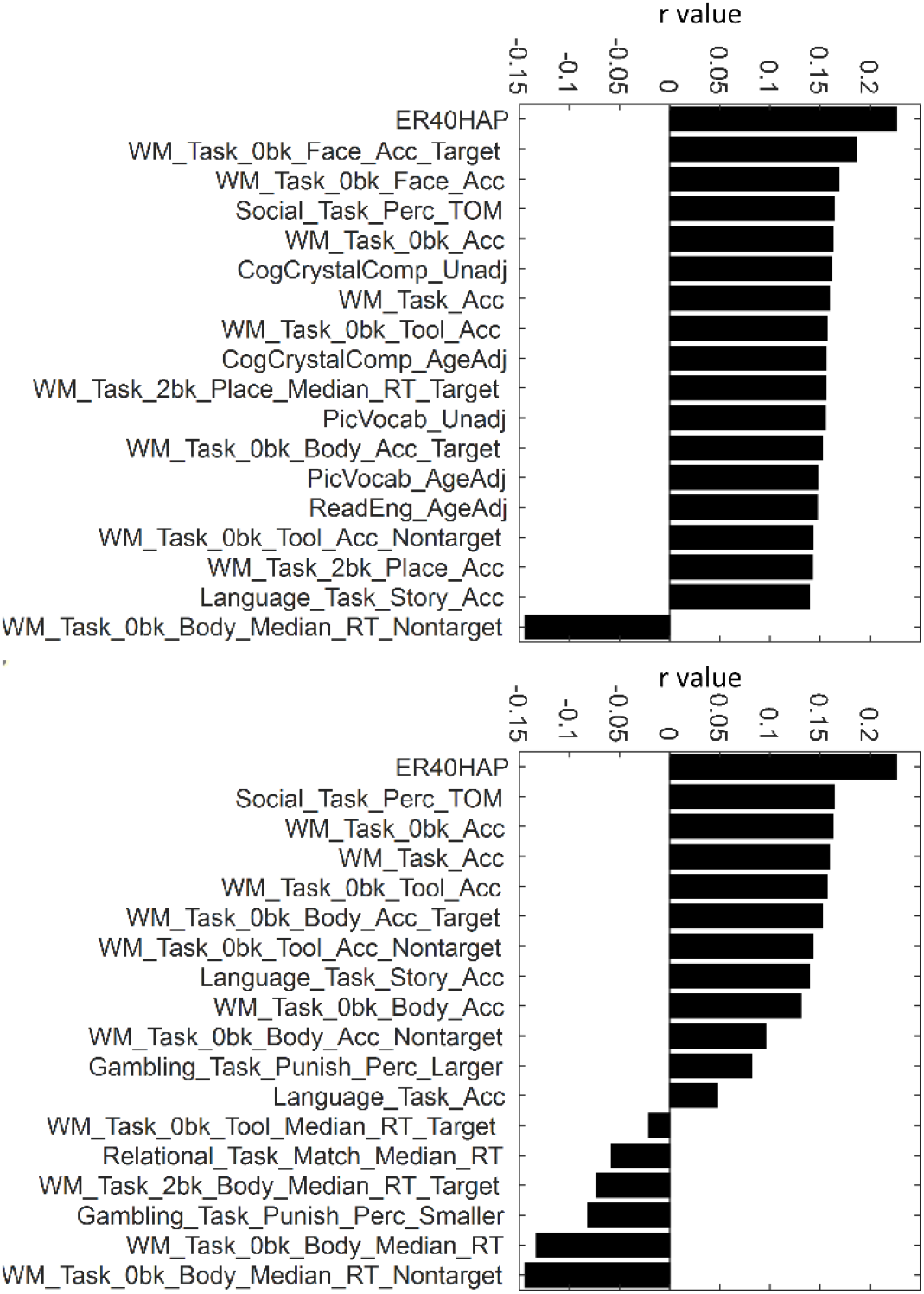
Correlation between the number of transitions in two major communities and behavior scores. The upper panel shows the correlations between the number of transitions in community 1 and behavior scores. The lower panel shows the correlations between the number of transitions in community 2 and behavior scores. Only significant correlations (p<0.05) are shown in this figure.

### The relationship between instantaneous HRFs and spectral patterns

As shown in Figure 8, the windowed HRFs vary with time as well, and their shapes are strongly coupled to the instantaneous spectral patterns, i.e., an HRF with decreased initial dip and increased time to peak is associated with an increased high-frequency component identified in power spectra. The five spectra modes correspond to distinct HRFs. The shifting peaks of modes 1-4 (shown in Figure 2) reflect the varying magnitudes of initial dips as well as time to peaks in HRFs. As mode 5 exhibits a spectral power relatively constant across the frequency band of interest, the initial dip and time to peak of its associated HRF appear to be the average those of modes 1-4.

**Figure 8.**
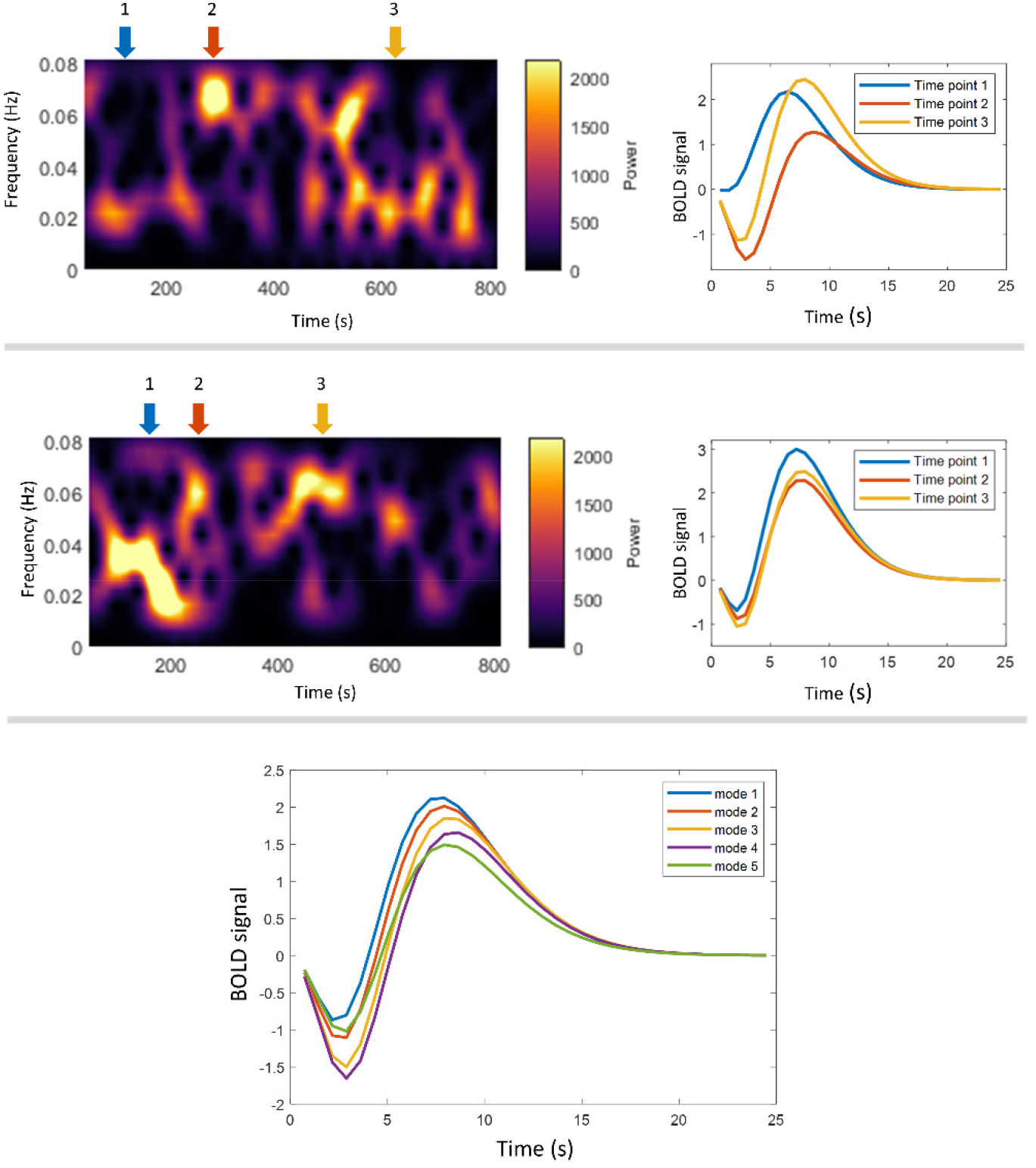
Relationship between instantaneous HRFs and spectral patterns. The upper and middle panel shows the HRFs (right) calculated in selected time windows where the spectral patterns (left) show peaks at different frequencies. Note that these two panels correspond to two voxels randomly selected from one subject. The lower panel shows the HRF corresponding to the five WM modes. Each line is the average HRF estimated from all time windows that have been assigned to a mode across time and voxels of the selected subject.

## Discussions

This study evaluated spectrograms of BOLD signals in WM in a resting state, which reflect the temporal variations of power spectral patterns within BOLD time courses. Applying k-means clustering to a set of measurements that consisted of such patterns at different time points and from different voxels and subjects, we obtained five dominant modes, equivalent to the centroids of the clusters. Each pattern was then assigned to a mode according to its distance to the centroids. Consequently, the original time course could be represented by the five modes as they varied over time. Several parameters, including occurrence, duration, and the number of transitions, were quantified and showed strong consistency across subjects. Besides that, one of the modes showed a strong coupling of its occurrence between GM and WM across subjects. We also identified two communities of voxels, where community 1 showed a higher probability of transitions among modes 1, 2, and 5 but a lower probability of transitions among modes 3, 4, and 5 than in community 2. Moreover, the total number of transitions in each community could predict specific human behaviors. Last, we observed that the five modes were associated with distinct HRFs.

Time-dependent frequency analysis is a well-established method for evaluating MEG and EEG data, and provides characteristic features of signal spectrograms which are associated with specific neurological conditions^27,28^. However, the frequency range of BOLD signals is much more limited so that spectrograms have rarely been applied in fMRI analyses. Recent studies have shown that the time-frequency pattern in GM is capable of describing the variabilities of functional connectivity in resting-state fMRI^29^ and could be categorized as a set of recurring modes^26^. These observations, along with our previous findings regarding the distinct power spectra identified in WM^25^, led us to investigate the spatial-temporal-frequency characterization of resting-state BOLD signals. The results inform us that the single/dual-peak power spectra calculated from the entire BOLD signal in our previous work^25^ are actually the unweighted combination of a series of time-varying spectral patterns. These patterns can be clustered into five modes, which occurred significantly more often at specific locations and whose distributions are consistent across subjects. However, we did not identify any voxel that showed significantly high occurrences of mode 2 across subjects. This partially explains the dual-peak patterns in power spectra (group level) that we observed in our previous study. That is, the lack of group consistency of the spatial distribution of the occurrence of mode 2 causes the trough (around 0.035 Hz) between the two peaks at a group level^25^. Moreover, three “high-frequency” modes that could be identified in WM were missing from GM. This is consistent with the single-peak pattern of power spectra that we observed in GM in our previous study^25^. In the current study, in GM, we identified only one unimodal mode, which is identical to WM mode 1 in shape and occurs significantly often in cortical areas that are more close to the surface. Our data also suggest that the occurrence of this mode in the cortex is positively associated with the occurrence of WM mode 1 across the entire WM. This finding suggests that the BOLD fluctuations of a specific band (centered at ≈ 0.02 Hz) reflect a global process.

The uniform mode, mode 5, exhibits substantially higher occurrence and duration than other modes, suggesting that BOLD fluctuations are maintained at a low power level across frequencies that uniformly vary between 0.01-0.08 Hz most of the time, accompanied by brief bursts (unimodal mode1 2,3,4) with higher power at more concentrated frequency ranges. A similar finding has been reported in GM in a previous study that investigated the dynamic functional connectivities among cortical areas^30^, suggesting that spontaneous brain activity may be dominated by brief periods of activity, possibly originating from a neuronal avalanche phenomenon. For unimodal modes, modes 1 and 2 occur significantly often in the inferior frontal WM, whereas modes 3 and 4 occur significantly often in the paraventricular and temporal areas. Moreover, these occurrences and durations are consistent across subjects, reflecting a highly reproducible pattern of WM BOLD signal, which has not previously been identified from the original time courses. The spatial distribution of the voxels where modes 3 and 4 frequently occur is similar to the areas with higher fiber complexity identified in our previous work^25^. Such crossing fiber bundles connect to multiple GM areas. Therefore, when these bundles received independent neural events from multiple GM they connected with at the same time or within small timing differences, they require more energy, lead to a higher level of oxygen extraction and consequently a lower initial dip we observed in the instantaneous HRFs. In addition, these bundles share the same blood supply, which, however, is insufficient to support the blood volume in processing the multiple events in a short time, causing the increased time to peak in HRF.

We identified two communities of voxels that showed distinct patterns of inter-mode transitions. Transitions between the uniform mode and unimodal modes were substantially more frequent than the transitions among unimodal modes. Particularly, direct transitions between modes 1, 2 and modes 3, 4 were rarely observed. These findings support the notion that mode 5 is a “baseline” mode from which a unimodal mode is likely to first transition before switching to other unimodal modes. The inferior frontal area exhibited a significantly lower number of transitions. We speculate that this is because activity linked to the inferior frontal area is suppressed during the resting-state scan, perhaps because it is associated with language and speech processing.

We observed that the occurrence and number of transitions of modes were significantly correlated with specific behavioral scores, though the correlation coefficients (r) were not high. However, the scores that significantly predicted cognitive abilities share some commonalities. For example, more than half of the scores shown in our data were positively associated with working memory accuracies. Meanwhile, negative correlations were mostly associated with the response time (RT) in working memory tasks. This observation is consistent with the notion that accuracies in working memory tasks are often negatively correlated with response times^31^. Future work is needed to address the relationship between these measurements and behaviors.

In summary, this is the first report of spatial-temporal-frequency characterization of BOLD resting-state time courses in white matter. Our results revealed the non-stationary nature of the spectral patterns of BOLD fluctuations in WM. The measurements of BOLD spectra were highly consistent and reproducible, including occurrence, duration, and transitions of modes, and reveal recurring patterns of power spectra as well as HRFs that have not been previously reported in WM BOLD signals, adding our understanding of spatial-temporal-frequency-physiological associations in the human brain.

## Methods

### HCP data

Two hundred subjects were randomly selected from the HCP S1200 release^32^. All were healthy young adults, 88 male and 112 female, whose ages ranged between 22 and 35 years. The imaging protocols are described in detail in a previous work^32^. Briefly, data were acquired using a 3T Siemens Skyra scanner (Siemens AG, Erlanger, Germany). The resting-state data were acquired using multiband gradient-echo echo-planar imaging (EPI). Each session consisted of two runs (left-to-right and right-to-left phase encoding) of 14 min and 33 s each, repetition time (TR) = 720 ms, echo time (TE) = 33.1 ms, voxel size = 2 mm isotropic, number of volumes = 1200. Physiological data, including cardiac and respiratory signals, were recorded during fMRI scanning. T1 images were acquired using a 3D magnetization-prepared rapid acquisition with gradient echo (MPRAGE), TR = 2400 ms, TE = 2.14 ms, voxels size = 0.7 mm isotropic. Assessments of cognitive ability in the HCP data include tasks from the University of Pennsylvania Computerized Neurocognitive Battery^33^ and the Blueprint for Neuroscience Research - funded NIH Toolbox for Assessment of Neurological and Behavioral function (http://www.nihtoolbox.org).

### Preprocessing

The images drawn from the HCP repository were preprocessed through the minimal preprocessing pipelines (MPP) as detailed elsewhere^34^. Briefly, T1 images were nonlinearly registered to MNI space using FNIRT^35^ and underwent a Freesurfer pipeline which produced surface and volume parcellations as well as morphometric measurements^36^. For fMRI, the pipeline included motion correction, distortion correction using reversed-phase encoding directions, and nonlinear registration to MNI space. We performed additional processing including regression of nuisance variables, including head movement parameters (using one of the outputs of motion correction in the MPP pipeline), and cardiac and respiratory noise modeled by the RETROICOR approach^37^, and followed by a correction for linear trends and temporal filtering with a band-pass filter (0.01 – 0.08 Hz). As the analyses were restricted to WM, a group-wise WM mask was reconstructed by averaging the WM parcellations that were derived from Freesurfer across all subjects and thresholded at 0.95. In a similar manner, a GM mask was also reconstructed but using a lower threshold due to higher individual variabilities therein. Specifically, 0.65 was selected as an optimal threshold by first setting a threshod of 0.5 which then was increased gradually with a step of 0.01 until there were no overlapping voxels between GM and WM. Finally, to increase the signal-to-noise ratio of each voxel, and to reduce the computation load, we downsampled the preprocessed images from 2mm resolution to 3mm.

### Calculation of the spectral modes

The workflow for calculating the spectral modes is shown in Figure 2. First, a spectrogram was calculated for the BOLD signal of each voxel using a short-time Fourier transform (STFT) ^38^ by first splitting the entire time course into a set of partially overlapped windows and calculating the transform for each window. Then the power spectrum of each window was calculated, showing the amplitude of BOLD fluctuations at different frequencies as they varied with time. In practice, we selected a window length of 100.4 s (140 TR) to provide a frequency resolution down to 0.01 Hz, which was considered the low cutoff of the baseline brain activity band^39^. To provide good temporal resolution while avoiding excessive computation load, we selected a step length of 2.88 s (4 TR). This resulted in 266 windows that overlapped by 97.52 s (136 TR) with each neighbor, evenly spanning the duration of the BOLD time course. By obtaining the spectrograms across voxels and subjects, we obtained a large number of observations (more than 490 million) of the spectral patterns across time. These observations were then grouped into a set of clusters using the K-means method. The elbow criterion^40^ was used to determine the optimal value of the number of clusters. Specifically, we successively increased the number of clusters from 1 to 20 and plotted the curve of the sum of the observation-to-centroid distances against the number of clusters. The optimal value of the number of clusters, which is five in our study, was determined by the location of the elbow on the curve. The five modes stand for the centroids of the clusters. Then each window-wise spectral pattern can be assigned to a specific mode according to its distance to the centroids.

### Calculation and statistics of the occurrence, stay time, and transitions of the modes

The original BOLD time course of each voxel can be represented by transitions between the five modes over time, including their frequency of occurrence and durations. The occurrences can be obtained by counting the numbers of each mode over the time course. The duration describes how long any mode persists before the signal switches to another mode. As each mode occurs multiple times during scanning, the duration is actually a mean time over multiple occurrences. A transition simply means a specific switch of modes from one to another. The total number of transitions is how many times we identify mode i at time point t and mode j at time t-1 where i ≠ j.

The occurrences, durations, and transition number can be measured for every voxel, resulting in maps of their distributions for each subject. Before group-level statistics, we performed Z transformations (i.e., we subtracted the global mean value and then divided by the standard deviation) to each map for each subject. The Z maps were spatially smoothed within the WM and GM masks separately with a 4 mm FWHM Gaussian kernel. Then a one-sample t-test was used to identify voxels that exhibited significantly higher or lower values compared to zero, the mean measurement across all voxels.

### Estimation of window-wise HRFs

HRFs were estimated from resting-state timecourses *b(t)* in each subject using a blind deconvolution approach^41,42^. The method requires no prior hypothesis about the HRF and is based on the notion that relatively large amplitude BOLD signal peaks represent the occurrence of separable, major, spontaneous events. In our study, first, such events were detected as peaks beyond a specified threshold (here, greater than 1.5 standard deviations over the mean). For each event, a general linear model was fitted using a combination of *s*_*n*_*(t)*, the onset of the event, and *h(t)*, which represents a linear combination of two gamma functions and its temporal derivative. Here *n* characterizes the time from the onset to the peaks *s(t)*, where *s(t)* =1 only if *t* corresponds to the peaks (events) we detected. The double gamma functions together with temporal derivative are capable of modeling an initial dip and time delay in the response ^43,44^. By searching for an *n (n∈ 0-12 s)* and minimizing the covariance of the residuals *cov[b(t) – conv(s*_n_*(t), h(t))]*, several parameters that model *h(t)* can be estimated, so the HRF *h*_*n*_*(t)* can be obtained.

## Acknowledgments

This work was supported by the National Institutes of Health (NIH) grant R01 NS093669 (J.C.G) and R01 NS113832 (J.C.G), and Vanderbilt Discovery Grant FF600670 (Gao). Imaging data were provided by the Human Connectome Project, WU-Minn Consortium (Principal Investigators: David Van Essen and Kamil Ugurbil; 1U54MH091657) funded by the 16 NIH Institutes and Centers that support the NIH Blueprint for Neuroscience Research; and by the McDonnell Center for Systems Neuroscience at Washington University.

We declare no conflict of interest

