## Supplement Figures for "Dynamic variations of resting-state BOLD signal spectra in white matter"

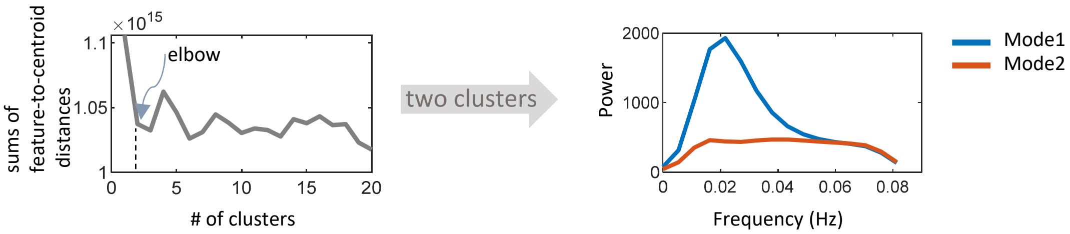


Figure S1. Spectra modes identified in GM. The workflow is the same as that for WM. Observations are grouped into two clusters using the K-means method. The elbow criterium is used to determine the optimal value of the # of clusters (2 in the current study).

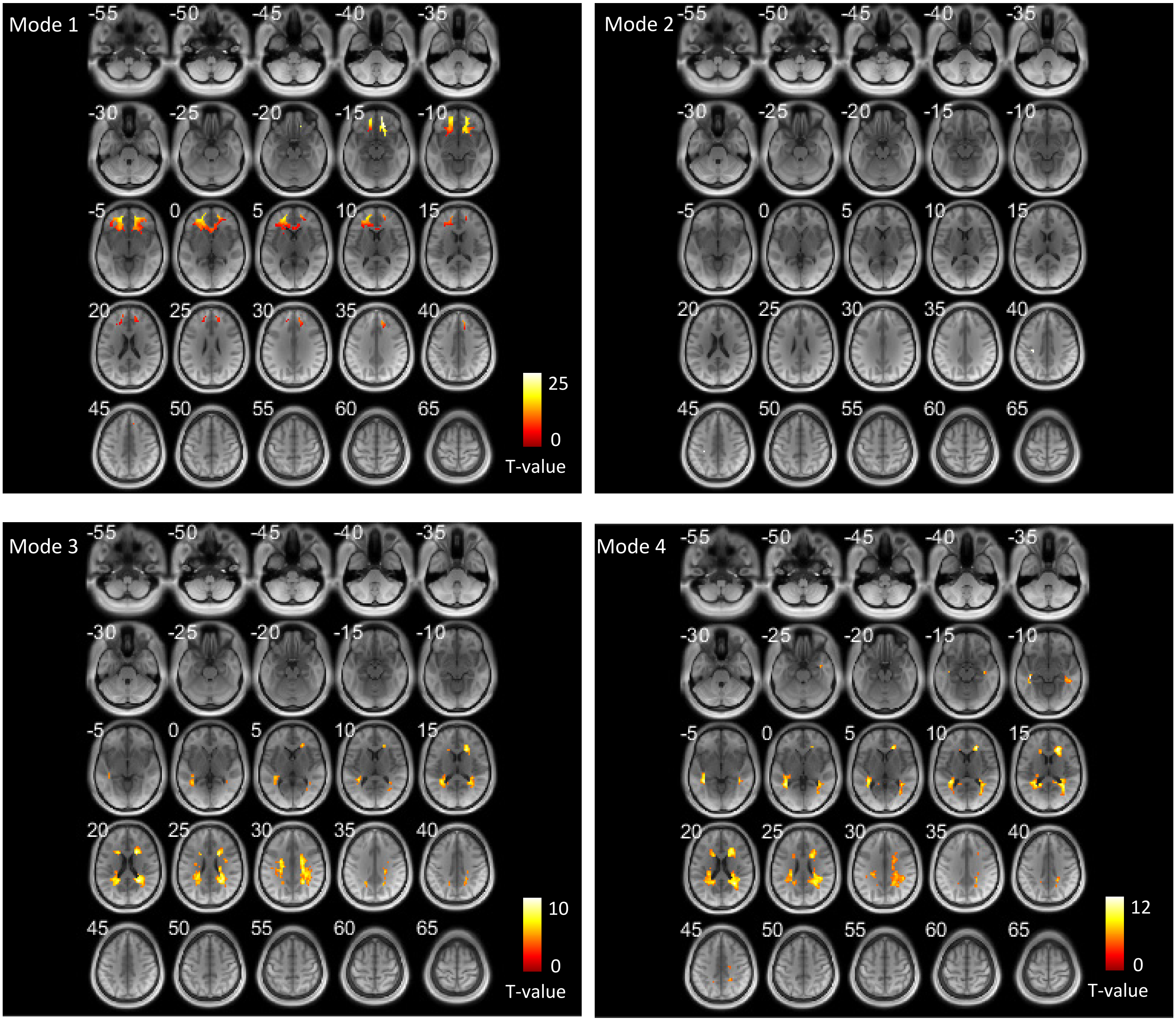


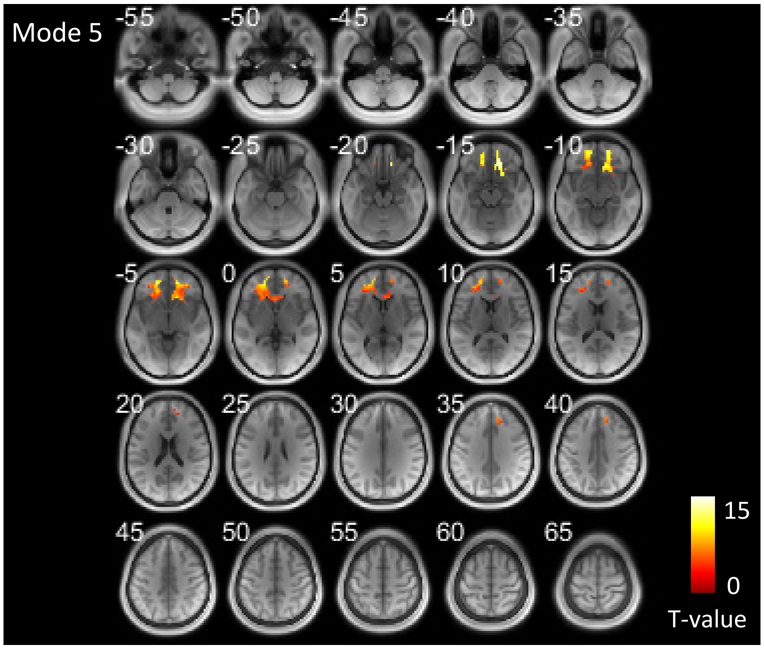


Figure S2. Areas that exhibit significantly high occurrence of the five modes in WM across all subjects shown in axial slices. (one-sample t-test, $p<0.05$ FWE corrected)

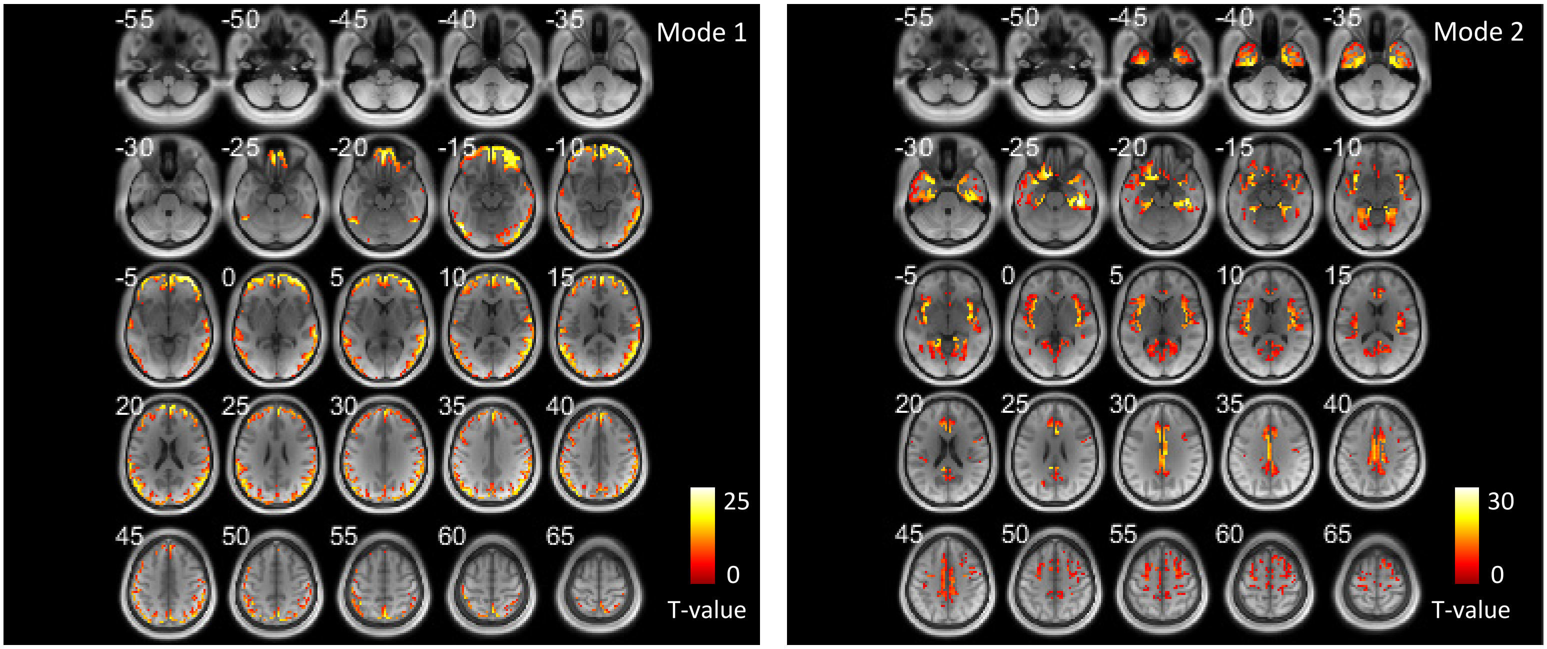


Figure S3. Areas that exhibit significantly high occurrence of the two modes in GM across all subjects shown in axial slices. (one-sample t-test, $p<0.05$ FWE corrected)

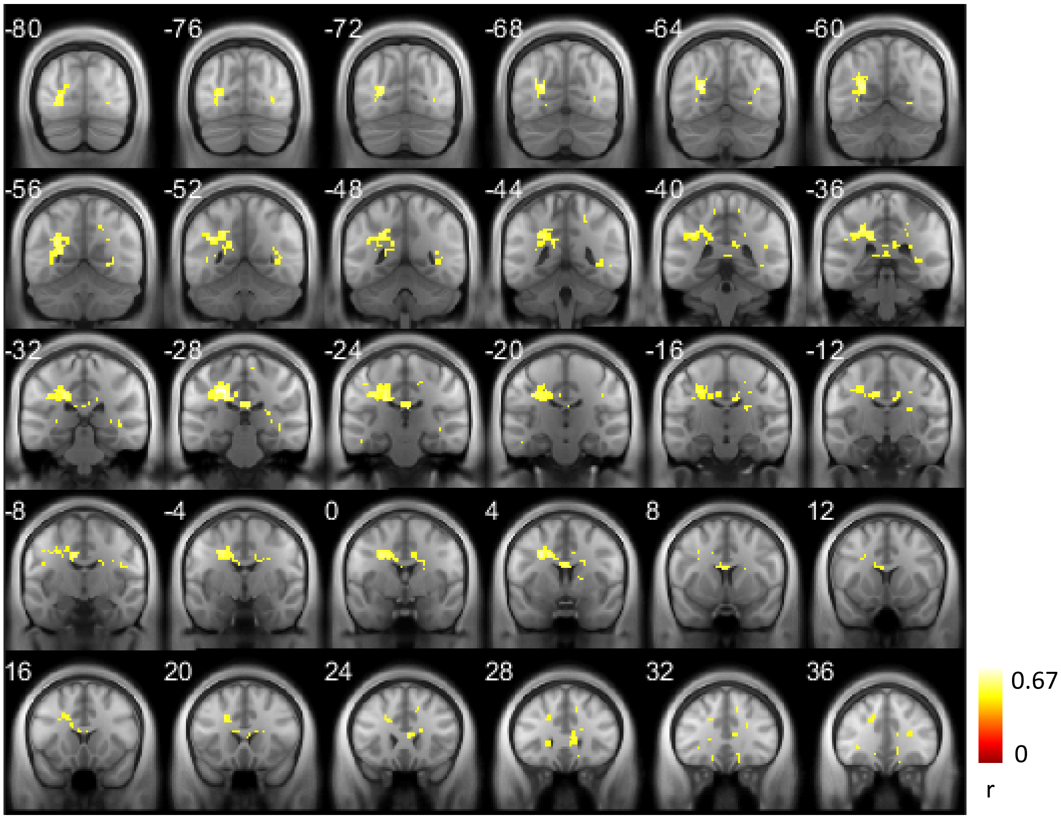


Figure S4. The coupling of mode occurrence (mode 1) in WM voxels to GM (mode 1) across subjects. (one-sample t-test, $p<0.05$ FWE corrected). This is a result beyond significance level corresponding to Figure 4 (left).

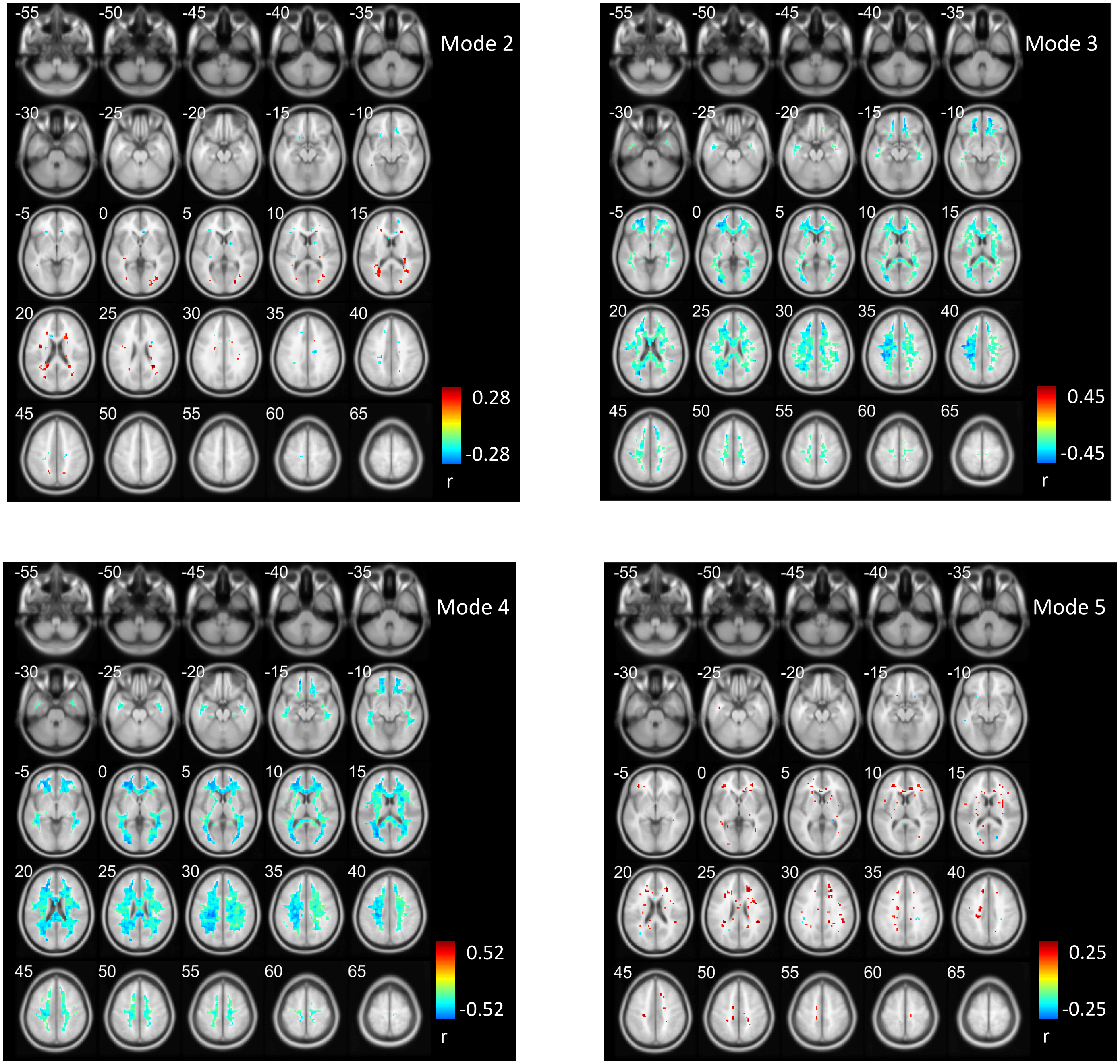


Figure S5. The coupling of mode occurrence (mode 2,3,4,5) in WM voxels to GM (mode 1) across subjects. (one-sample t-test, $p<0.05$ FWE corrected)

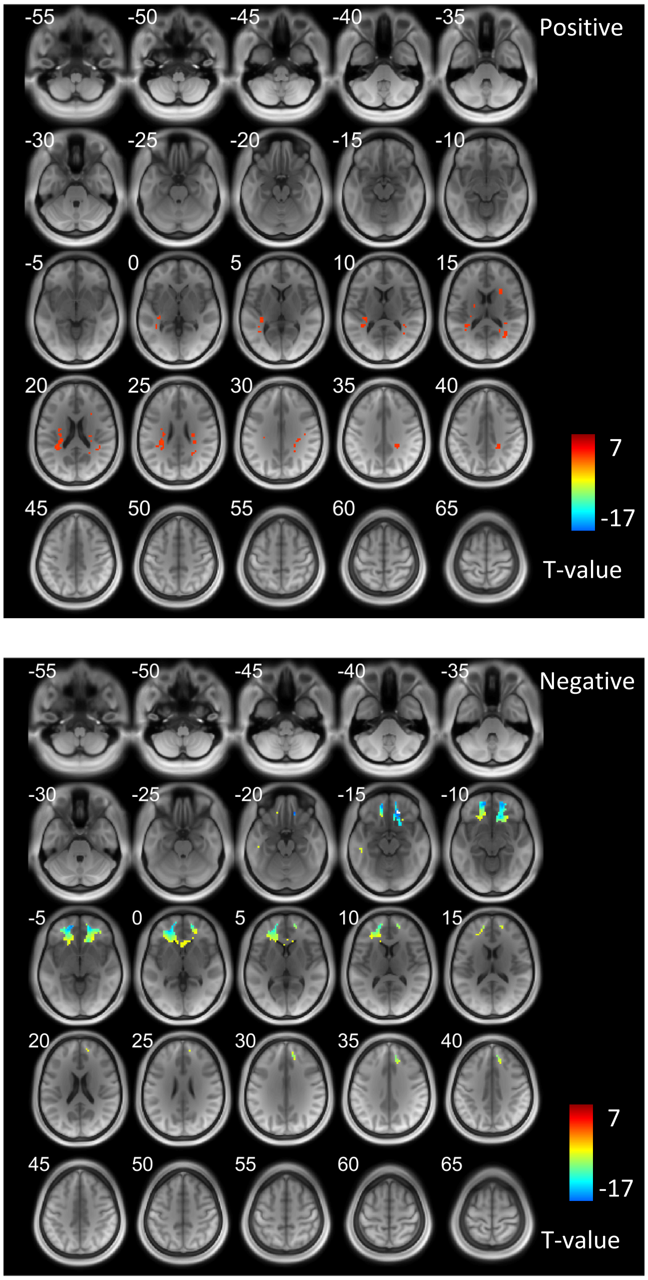


Figure S6. The number of transitions of each WM voxel across all subjects. (one-sample t-test, p<0.05 FWE corrected). This is the result beyond significance level corresponding to Figure 5.

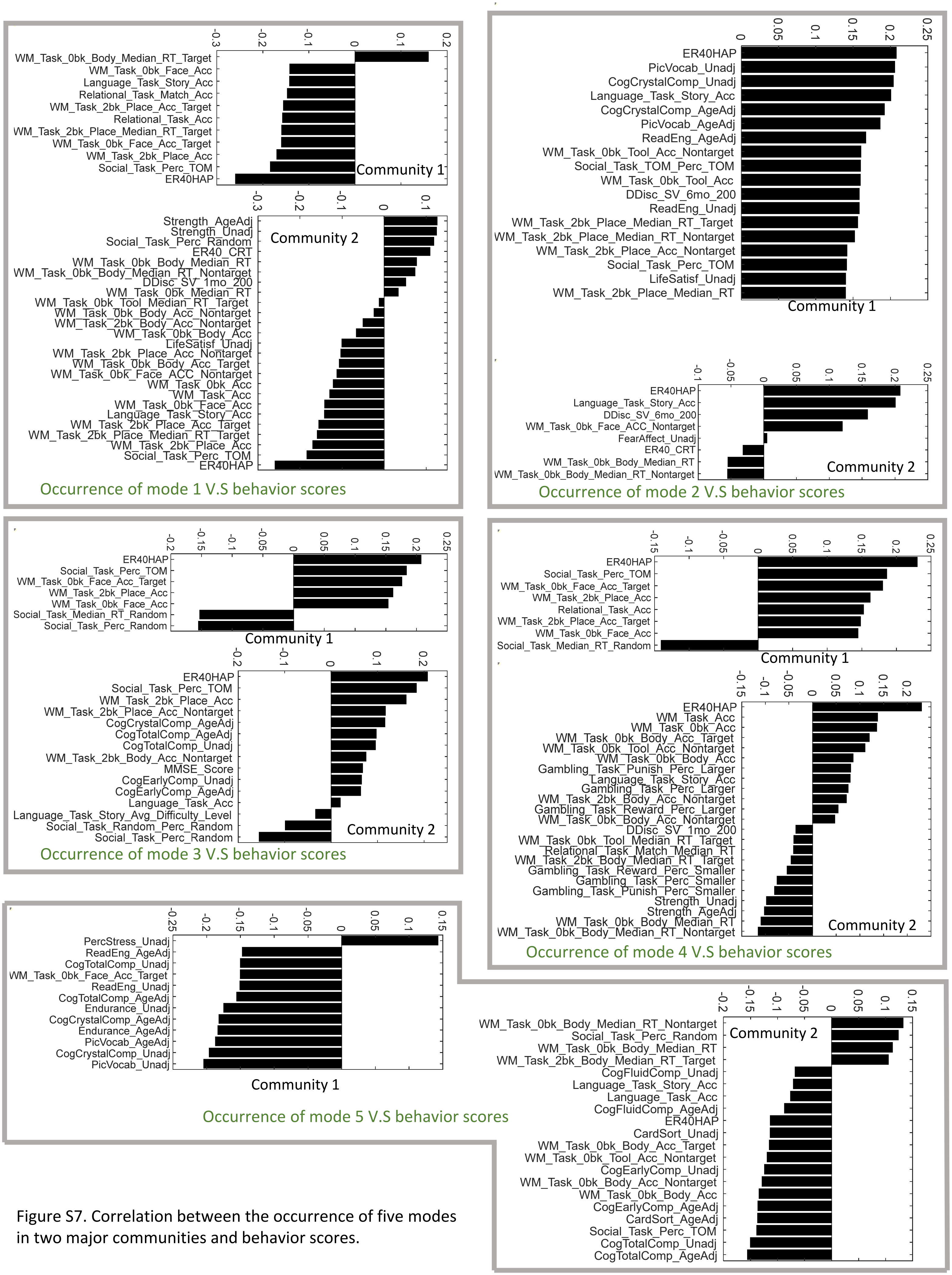


Figure S7. Correlation between the occurrence of five modes in two major communities and behavior scores.
